# NeuroDecodeR: A package for neural decoding analyses in R

**DOI:** 10.1101/2022.12.17.520811

**Authors:** Ethan M. Meyers

## Abstract

Neural decoding is a powerful method to analyze neural activity. However, the code needed to run a decoding analysis can be complex, which can present a barrier to using the method. In this paper we introduce a package that makes it easy to perform decoding analyses in the R programing language. We describe how the package is designed in a modular fashion which allows researchers to easily implement a range of different analyses. We also discuss how to format data to be able to use the package, and we give two examples of how to use the package to analyze real data. We believe that this package, combined with the rich data analysis ecosystem in R, will make it significantly easier for researchers to create reproducible decoding analyses, which should help increase the pace of neuroscience discoveries.

## Introduction

Neural decoding is a data analysis method that uses neural activity to predict which experimental conditions are present on different experimental trials (E. M. Meyers & Kreiman, 2012; Quian Quiroga & Panzeri, 2009). Neural decoding analyses have been used on data from a range of recording modalities in humans, including on electroencephalogram (EEG) and magnetoencephalogram (MEG) signals (Carlson et al., 2011; Isik et al., 2014), electrocorticography (ECoG) recordings (Volkova et al., 2019), single unit recordings (Rutishauser et al., 2015; Saha et al., 2021), and fMRI BOLD responses, where the method is referred to as multivariate pattern analysis (Haynes & Rees, 2006; O’Toole et al., 2007; Pereira et al., 2009; Tong & Pratte, 2012; Weaverdyck et al., 2020). Neural decoding analyses of spiking activity have also been conducted in a range of animal species and in different brain regions including the motor cortex of macaques to predict reaching directions and control brain computer interfaces (Georgopoulos et al., 1986; Wessberg et al., 2000), the hippocampus of rats to predict spatial location information (Brown et al., 1998; Tingley & Buzsáki, 2018), the inferior temporal cortex of macaques to predict which visual objects were present (Hung et al., 2005; Zhang et al., 2011), and higher level brain regions to predict a range of cognitive related variables (Crowe et al., 2010; Rikhye et al., 2018).

There are several advantages to using neural decoding to analyze data including the ability to pool signals across many recording channels which can give a clearer picture of what information is in a brain region at a particular point in time (Quiroga et al., 2004). Additionally, neural decoding can be used to assess how information is coded in populations of neural activity (Jacobs et al., 2009; E. M. Meyers et al., 2015; Nirenberg & Latham, 2003), such as whether there is a small subset of neurons that contain all the information present in a larger population (E. M. Meyers et al., 2008a, 2012) and whether information is coded by patterns of activity that change in time (King & Dehaene, 2014; Meyers, 2018; Meyers et al., 2008a).

Despite the advantages neural decoding has as a data analysis method, the code needed to run a decoding analysis can be complex which can present a barrier to using the method. To address this difficulty, several software packages exist that make it easier to run these analyses including packages in Python (Glaser et al., 2020; Hanke et al., 2009) and MATLAB (Hebart et al., 2015; Oosterhof et al., 2016; Peng et al., 2020). In previous work we have also tried to address this issue by creating a MATLAB toolbox called the Neural Decoding Toolbox (Meyers, 2013).

In this paper, we introduce a new neural decoding package written in R (R Core Team, 2021), called NeuroDecodeR. The design of the NeuroDecodeR is based on the design of the MALTAB Neural Decoding Toolbox, but it extends its functionality several ways. In particular, the NeuroDecodeR includes the ability to easily add new measures for quantifying decoding accuracy, a system to manage results, and the ability to run the code in parallel which greatly speeds up the time it takes to run an analysis. Additionally, using the R programming language has several advantages including that R is free/open source, and that there is a large data analysis ecosystem for creating reproducible data analyses.

In the following paper, we describe the design of the NeuroDecodeR package, the data format that is used by the package, and we give an example of how the package can be used by reproducing results from Meyers et al (2015). We hope this package will make it easier for neuroscientists to extract insights from the data they collect, and will help introduce neural decoding analyses to the larger Statistics/Data Science community that uses R to analyze data.

## Methods

### A brief overview of neural population decoding

Neural decoding is a data analysis method that assesses whether information about particular stimuli, or other behaviorally relevant variables, is present in neural activity (Quian Quiroga & Panzeri, 2009). The method works by ‘training’ a machine learning algorithm, called a *pattern classifier*, to learn the relationship between neural activity and particular experimental conditions on a subset of data called the *training set*. Once the classifier has ‘learned’ the relationship between the neural data and experimental conditions, one assesses whether this relationship is reliable by having the pattern classifier predict which experimental conditions are present in a separate *test set* of data.

In a typical analysis, the data in split into k different parts, and the classifier is trained on k-1 parts and tested on the remaining part. This procedure is repeated k times where a different part of the data is used to test the classifier each time, in a process called *cross-validation*, and a final measure of prediction accuracy is aggregated across the performance on all k test sets. If the pattern classifier can make accurate predictions on these separate test sets of data, then this indicates that a brain region has information about a particular experimental condition (for more information about decoding see Meyers & Kreiman, 2012).

Decoding methods are often applied to time series data, such as neural activity recorded over a fixed length experimental trial. To do this, the classifier is trained and tested at one point in time, and then the procedure is repeated at the next point in time. This leads to results that show how the information content fluctuates over the course of a trial (for an example, see Figure 3A), and can be used to assess how information flows through different brain regions (Meyers et al., 2018). Additionally, neural decoding can be used to gain insight into how information is coded in neural activity. For example, a temporal cross-decoding (TCD) analysis can be done where the classifier is trained at one time period and then tested at a different time period (for an example, see Figure 3B). If the classifier has a high decoding accuracy when trained and tested at the same time period, but a low decoding accuracy when trained and tested at different time periods, then this indicates information is coded by patterns of activity that change in time (King & Dehaene, 2014; Meyers et al., 2008b). Abstract information can also be assessed using a ‘*generalization analysis’* by training the classifier on one set of conditions and then testing the classifier on a related set of conditions. For example, Hung et al., (2005) assessed whether the inferior temporal cortex contains information about objects that is invariant to the position by training the classifier to discriminate between a set of objects that were shown at one retinal position and then testing the classifier to see whether it could make predictions at a different retinal position (also see “Decoding analysis 2” below for another example of a generalization analysis).

When running a decoding analysis, neural activity from different sites do not need to be recorded simultaneously, but instead one can create ‘pseudo-populations’ where responses of simultaneously recorded neural populations are approximated by combining recordings made across multiple experimental sessions (Averbeck et al., 2006; Meyers & Kreiman, 2012). These pseudo-populations allow a larger number of sites to be included in a decoding analysis, which can lead to clearer results. As described below, one of the strengths of the NeuroDecodeR package is that it can automatically create pseudo-populations as part of the decoding procedure.

### Installing the NeuroDecodeR package

The NeuroDecodeR package has been published on the comprehensive R archive (CRAN). Like all R packages, the NeuroDecodeR package must be installed before it can be used for the first time. To install the package, use the command: `install.packages(“NeuroDecodeR”)`. Once the package is installed, any time you would like to use it you can load it into memory using: `library(NeuroDecodeR)`.

Online documentation for the NeuroDecodeR package is available at https://emeyers.github.io/NeuroDecodeR/. The development version of the package is available on GitHub at https://github.com/emeyers/NeuroDecodeR.

### Design of the NeuroDecodeR package

The NeuroDecodeR package is designed around five abstract object types which allows researchers to easily experiment with a range of different decoding analyses. The five object types are:

1. <u>Datasources (ds):</u> These objects create training and test splits of the data.
2. <u>Feature preprocessors (fp):</u> These objects extract statistics from the training set, and then apply transformations to the training and test set. These transformations are typically used to either improve the decoding accuracy, or to assess how information is coded in neural activity.
3. <u>Classifiers (cl):</u> These objects learn the relationship between neural activity and experimental conditions on the training set of data, and then make predictions on the test set of data.
4. <u>Result metrics (rm):</u> These objects compare the predictions made by the classifier on the test set of data to the experimental conditions that were actually present and then create metrics that indicate how accurate the predictions were. The final decoding results are stored in these objects as data frames and can be plotted using associated plot methods.
5. <u>Cross-validator (cv):</u> These objects take a datasource, feature preprocessors, a classifier, and result metrics, and run a decoding analysis by:
  a. Generating training and test data from the data source.
  b. Pre-processing the data using the feature preprocessors.
  c. Passing the data to the classifier which learns a model on the training data and then makes predictions on the test data.
  d. Passing the classifier’s predictions to the result metrics which creates measures of how accurately the information can be decoded.
  Steps a-d are typically repeated over several times on different “resample runs” where different training and test sets are created on each run which can lead to more accurate results.

The NeuroDecodeR package comes with one or more implementations of each of these object types, as listed in Table 1. A description of how to use these objects to run a decoding analysis is shown in the Results section below.

**Table 1.**
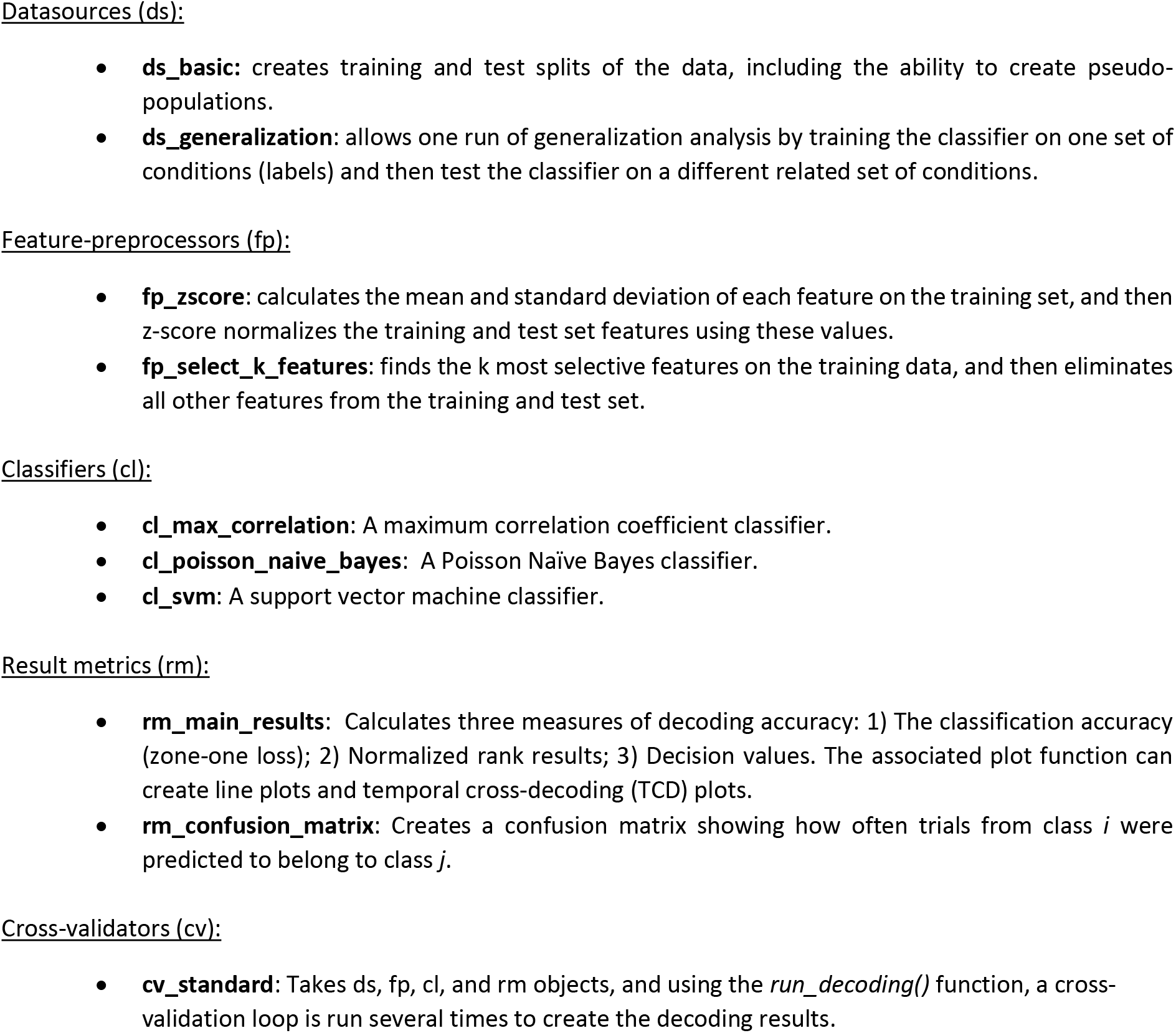
A list of implementations NeuroDecodeR objects that come with the NeuroDecodeR package. The package also includes a number of additional functions that are useful for processing data, plotting, and saving/loading results. For more details on these objects, see Supplemental Table 1, and the online documentation at: https://emeyers.github.io/NeuroDecodeR/reference/index.html

The advantage of this modular design is that it allows researchers to easily try out different analyses to gain additional insights and to make sure the results are robust to particular analysis choices. For example, as described below in the Results section, one can use the `ds_generalization` datasource instead of the `ds_basic datasource` to run a generalization analysis which can assess whether there is abstract information in the neural data.

Each of these object types are defined by an interface which specifies exactly which methods each object must have to work with the other object types in the package. This design allows users to add new implementations of the objects to their analysis. For example, a researcher could create a new classifier by implementing an S3 object that has a `get_predictions()` method.^1^ More information on how to implement the methods needed to create new NeuroDecodeR objects is available on the NeuroDecodeR’s documentation: https://emeyers.github.io/NeuroDecodeR/articles/NDR_object_specification.html.

### Data formats

#### Raster format

In order to use the NeuroDecodeR package, neural data must be put into a particular format called “raster format”. In this format, data from each recording site is in a separate “raster data” file. The data in each of these raster data files consists of a data frame where each row corresponds to one experimental trial, and each column must start with the prefix `site_info., labels.`, or `time.`. Columns that start with the prefix `site_info`. contain meta-information about the recording site. Columns that start with the prefix labels. contain information about which experimental conditions were present. Columns that start with the prefix `time`. contain the neural activity that occurred at a particular point in time.

To illustrate the data formats NeuroDecodeR package uses, and how to run a decoding analyses, we will use data from the “Friewald Tsao Face Views AM data set” (Freiwald & Tsao, 2010). This data set consists of recordings made from neurons in the macaque anterior medial face patch (AM) while monkeys viewed a random sequence of images where each image was presented for 200 ms with a 200 ms inter-stimulus interval. The images shown consisted of faces from 25 different people taken from 8 different head orientations^2^. For readers who are interested in replicating the analyses described below, a zip file that contains a directory with raster data from this experiment can be downloaded from http://www.readout.info/downloads/datasets/freiwald-tsao-face-views-am-dataset/. When this data archive is unzipped, a directory will be created called *Freiwald_Tsao_faceviews_AM_data*/ that has 193 files of data in raster format that we will use in the analyses.

Figure 1A shows data from one neuron that is in raster format. From looking at the raster format data we see the that the “site_info” columns consists of `site_info.monkey` which contain information about the name of the monkey and `site_info.region` which contains information about the brain region where the recording was made. We also have “labels” columns that list what stimulus occurred on each experimental trial including `labels.person` which indicates who the person was in each image that was presented, `labels.orientation` which indicate the head orientation in each image, and `labels.orient_person_combo` which combines the head orientation and person information. Finally, we have the “time” columns which contain the recorded data, including `time.2_3`, which contains the neural activity that occurred in the time window [2 3) milliseconds after the stimulus onset. Since the recordings are neuronal spiking activity, each data value in these “time” columns is either a 1 indicating a neuron produced an action potential or a 0 indicating that it did not. As described below, we can visualize raster format data using the `plot(raster_data)` function, which will produce plots such as the one shown in Figure 1B.

**Figure 1:**
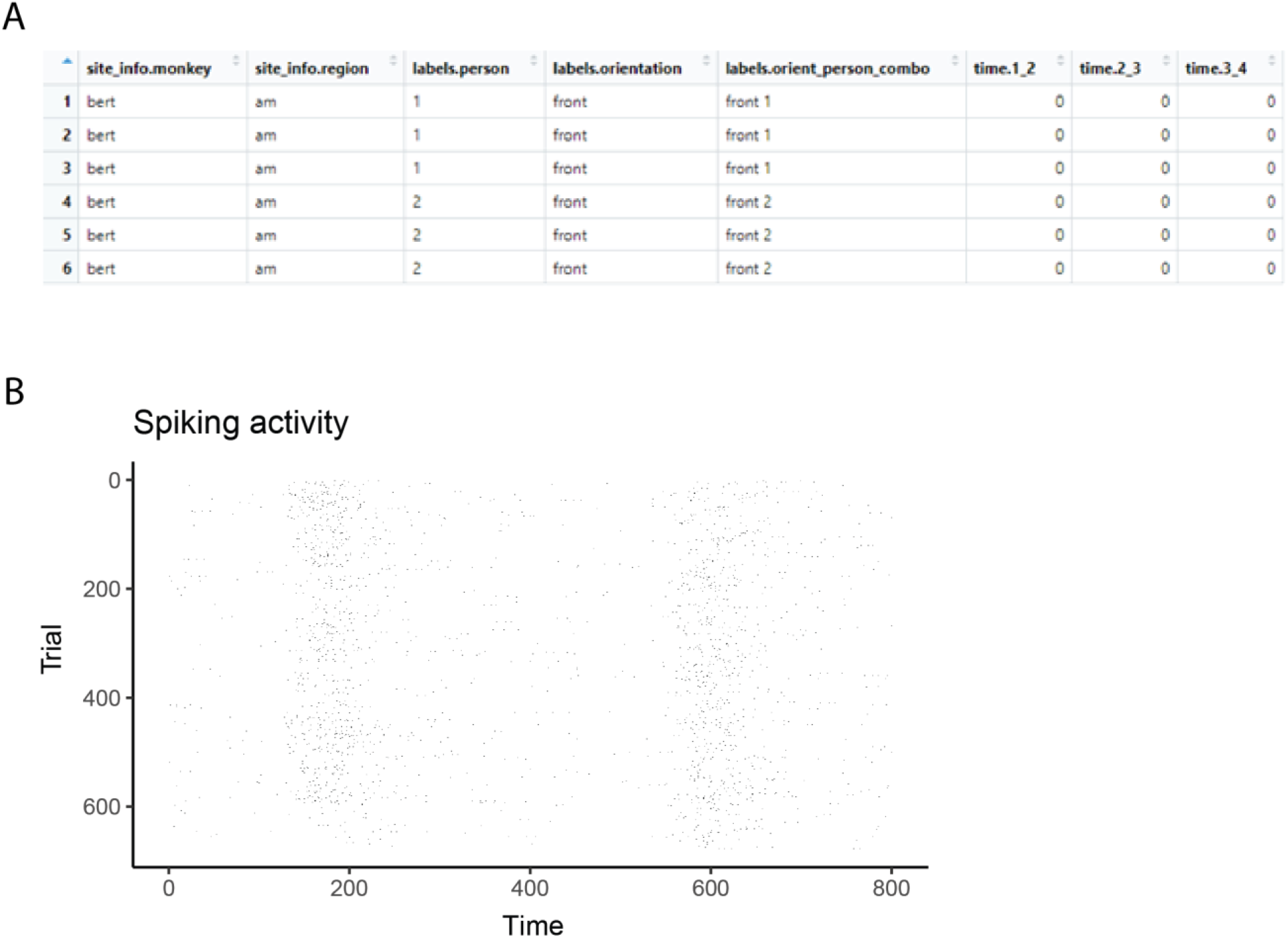
Stimuli and raster data from the Friewald Tsao Face Views AM data set. **A**. Example of the 8 head orientation stimuli from 1 of the 25 individuals in the Friewald and Tsao Face Views data set (the full data set consists of 25 individuals from these 8 head orientations). The labels below each image correspond the `labels.orientation` column in the raster data. **B**. An example of data in raster format. Each row corresponds to an experimental trial, and the columns start with either `site_info., labels`., or `time`. which is required for data to be in raster format. **C**. A visualization of data that is in raster format created by using the `plot(raster_data)` function.

When analyzing data from a new experiment, one can put data into raster format by saving data in comma separated value (csv) files with the appropriate column names (i.e., columns names that start with `site_info., labels.` and `time.`). The data can then later be loaded in raster format using the read_raster_data() function. Alternatively, one can save them in the MATLAB Neural Decoding Toolbox raster format (http://www.readout.info/toolbox-design/data-formats/raster-format/) and then convert the files into R using the `convert_matlab_raster_data()` function.

#### Binned format

Once one has created a directory that has data files from each recording site in raster format, one can convert this data to “binned format” using the `create_binned_data()` function. After this conversion is done, the rest of decoding process relies only on data in binned format. Data in “binned format” is similar to data in raster format in that it contains the same `site_info., labels`. and `time`. columns.; however, the binned data time. columns contain data at a coarser temporal resolution that is created by averaging activity in sliding time windows. For example, the Friewald Tsao Face Views raster data is recorded at millisecond resolution, as indicated by the fact that the time bins are successive numbers; i.e., *time.1_2, time.2_3, time.3_4*, etc. However, as described in the Results section, we can use the `create_binned_data()` to create averaged firing rates in 30 ms bins sampled every 10 ms, which will give us time bins columns with values `time.1_31, time.11_41, time.21_51`, etc. Additionally, data in binned format contains data from all recording sites as rows in the data table, and there is a `siteID` column that indicates which recording site the data in each row came from. The reason for binning the data is that it is usually computationally too expensive in terms of memory and runtime to analyze data at a very high temporal resolution. Additionally, firing rates averaged over longer time periods usually lead to higher decoding accuracies which can give clearer results (Meyers et al., 2009).

## Results

In the following sections, we describe how to use the NeuroDecodeR package to run two different decoding analyses. The analyses described below can be replicated by downloading the raster data at http://www.readout.info/downloads/datasets/freiwald-tsao-face-views-am-dataset/.

### Viewing raster data

As mentioned above, to illustrate how to use the NeuroDecodeR package we will analyze the “Friewald Tsao Face Views AM data set” which consists of 193 files in raster format that are stored in a directory called Freiwald_Tsao_faceviews_AM_data/. To begin we will load one of these raster format files using the `read_raster_data()` function, and then we can plot the data using the `plot()` function as follows:

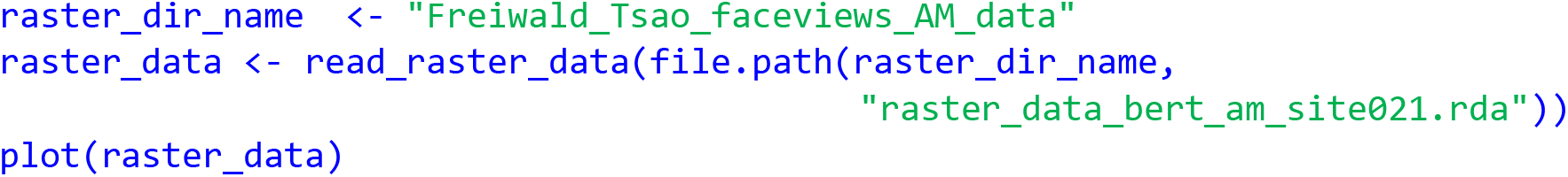

The results from visualizing the raster data for this neuron are shown in Figure 1B. In the plot, the x-axis corresponds to time from the stimulus onset, y-axis corresponds to different experimental trials, and black tick marks on the plot corresponds to the time when action potentials occurred.

While one does not need to visualize raster data to run a decoding analysis, it can be useful to visually examine raster data from a few sites to make sure the data conform to expectations. For example, in the Freiwald and Tsao experiment, images were shown every 400 ms (i.e., images were presented for 200 ms followed by a 200 ms interstimulus interval). When looking at the plot of the raster data example neuron shown in Figure 1B, we see that there is a large increase in spiking activity a little before 200 ms post stimulus onset, which corresponds to the response latency of this neuron, and then another large increase in spiking activity a little before 600 ms post stimulus onset, which corresponds to the response of the next stimulus. Seeing that the pattern of responses matches what we expect based on the design of the experiment is a good sanity check that we have correctly formatted the data.

### Binning the data

As we also mentioned above, all decoding analyses use data that is in binned format which can be created from raster data files using the `create_binned_data()` function. The `create_binned_data()` takes the following arguments:

1. A string specifying the path to a directory that contains the raster data files.
2. A string specifying a prefix that will be appended to the saved binned data file name.
3. A number specifying the bin width which neural activity will be averaged over.
4. A number specifying the sampling interval indicating the frequency with which to create repeat the binning process^3^.

For our analysis, we will average the activity in 30 ms bins, and we will sample the data at 10 ms intervals. This can be achieved by running the following command:

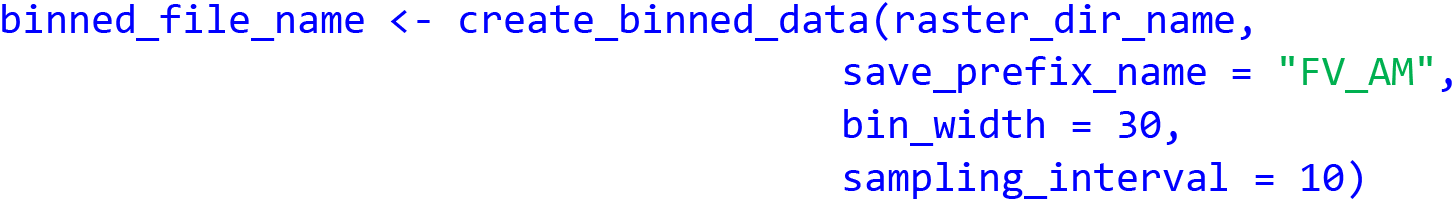

The resulting binned data is in a file called `FV_AM_30bins_10sampled.Rda` which we will use for all the subsequent decoding analyses in this paper.

#### Decoding analysis 1: Decoding face identity using left profile images

As a first demonstration for how to run a decoding analysis, we will decode which of the 25 individuals were shown on each trial using *only the left profile images*. To do this, we will use the `labels.orient_person_combo` column of the binned data, and we will only use the label levels that start with `“left profile”` which correspond to trials when left profile images were shown; i.e., we will use the label levels `“left profile 1”, “left profile 2”`, up to `“left profile 25”`, which corresponds to a trials where a left profile image of person 1 was shown, up to trials when a left profile image of person 25 was shown, and we are excluding trials when faces of other orientations were shown such as `“right profile 1”`, and `“frontal 10”`, etc.

### Assessing how many recording sites and cross-validation splits to use

Before starting to run a decoding analysis, it is useful to assess how many trials were collected from each recording site for each stimulus that was shown; for example, for neuron with siteID 7, how many trials were recorded when the image `“left profile 3”` was shown, etc. By examining this information across all the sites that were recorded, one can assess how many cross-validations splits to use, and whether specific sites should be excluded from further analyses because not enough trials were recorded from a given site. This is particularly important when creating pseudo-populations from experiments where data from different sites were recorded in different experimental sessions, since it is likely that sites that were recorded in different sessions will have different numbers of trials for each stimulus. If one uses k cross-validation splits, then only sites that have at least k trial repetitions *of all the stimuli* can be included in the analysis. Therefore, finding the sites that have enough repetitions of all the stimuli is an important first step so that one can tell which sites have enough data to be included in the analysis.

To visualize how many sites have at least k repetitions of each stimulus for different values of k, we can use the `get_num_label_repetitions()` function, along with the associated `plot()` function:

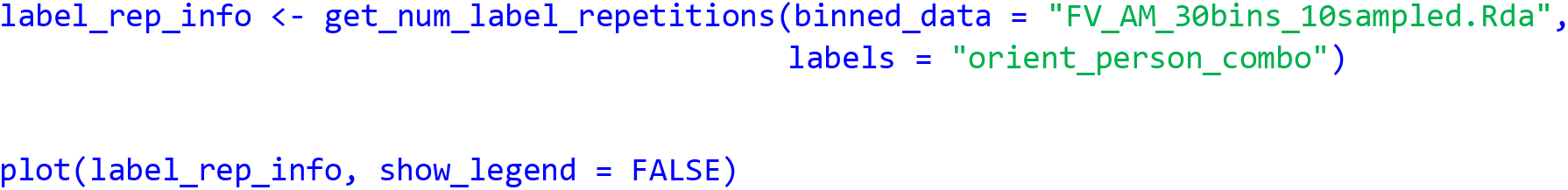

The results from running this code, shown in Figure 2, illustrate the trade-off between the number of cross-validation splits we would like to use (k), and the number of neurons available. To interpret this plot, we will focus on the black dashed line which shows how many sites have k repetitions of *all* the stimuli. From looking at this black dashed line, we see that there are a little less than 150 neurons that have 3 repetitions of each of the 25 stimulus, and there are a little more than 75 neurons that have 4 repetitions of each stimulus. Thus, if we run a 3 fold cross-validation analysis our pseudo-population vectors could consist of a little less than 150 neurons and if we run a 4 fold cross-validation analysis, our pseudo-population vectors could consist of a little more than 75 neurons. In the subsequent analyses, we will run a 3 fold cross-validation, although a 4 fold cross-validation with fewer neurons would also be a reasonable choice^4^.

**Figure 2:**
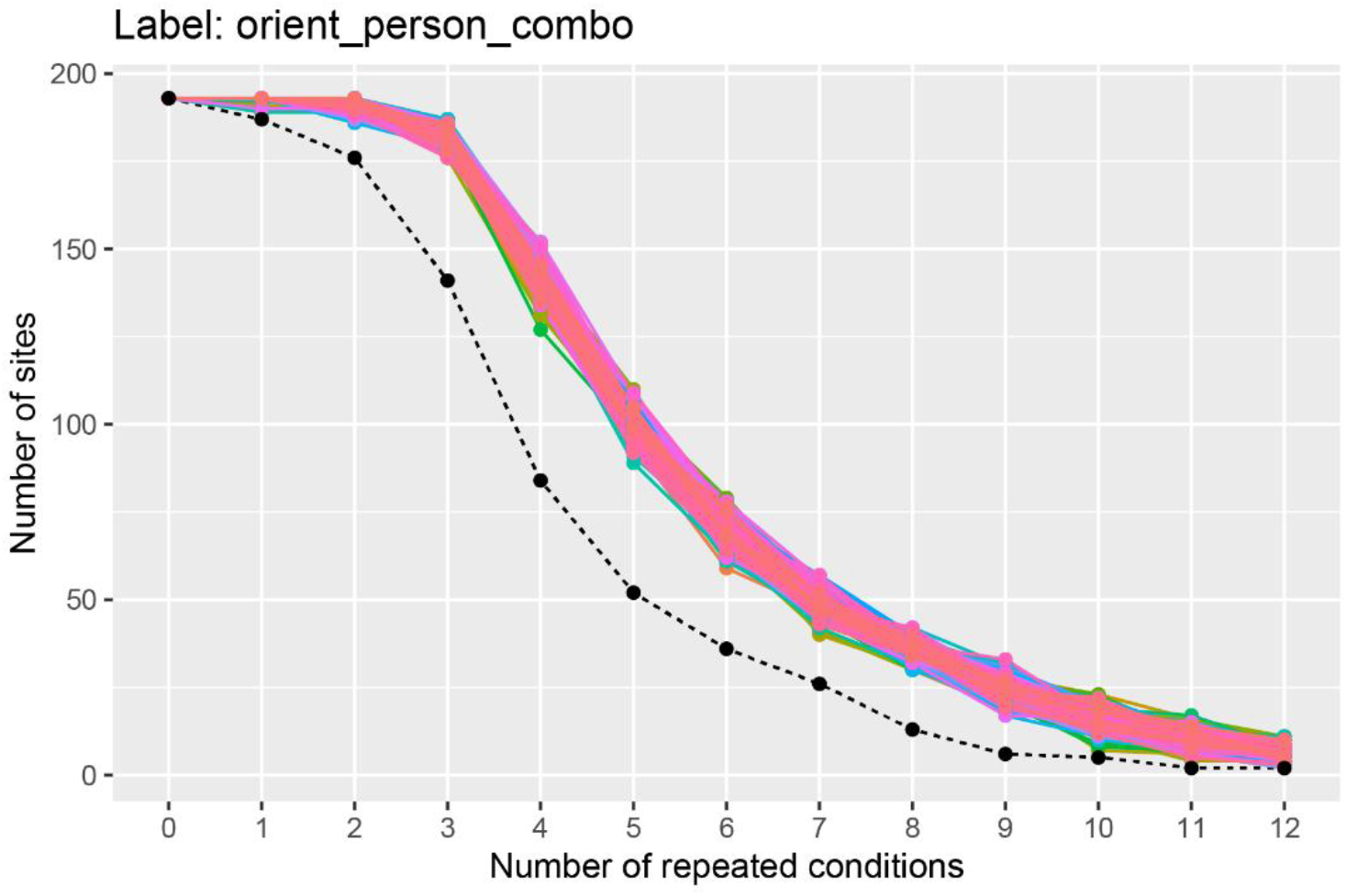
A plot showing how many sites (i.e., neurons) have at least k repetitions of all label levels for the orient_person_combo label. The black dashed line shows how many sites have at least k repetitions of for *all* label levels, and the colored traces show how many sites have at least k repetitions for each specific label level in the data set; since there are 8 * 25 = 200 label levels (i.e., stimuli), there are 200 colored lines on this plot. As expected, as the number of repeated conditions k increases, there are fewer sites available that have k repetitions of all the label levels. When running a decoding analysis, one needs to select a number of cross-validation splits k, and only sites that have at least k repetitions of all label levels can be used in the analysis. Thus, there is a trade-off between how many cross-validation splits to use and how many sites are available for decoding. The black dashed line is useful for selecting the number cross-validations splits k to use so that one has both a reasonable number of cross-validation splits and a reasonable number of sites available. Additionally, the colored lines are useful for assessing if particular label level (e.g., stimulus) have far fewer repetitions which would be the case if a particular colored line was much closer to the black dashed line than the other colored lines (which is not the case here). If a particular label level has far fewer repetitions than other label levels, then one can exclude the label level from the analysis in order to increase the number of sites available for decoding.

The colored lines on the plot show how many sites have at least k repetitions for each specific stimulus, where there is a different colored line corresponding to each stimulus that was shown in the experiment. If one of these colored lines was close to the black dashed line, this would indicate that there was a stimulus that had fewer repetitions than the other stimuli and we might consider excluding this stimulus from our analysis to make more sites available to use in our analysis.

To restrict our subsequent analyses to only use sites that have 3 repetitions of all stimuli, we can use the `get_siteIDs_with_k_label_repetitions()` functions which will give us the site IDs of all neurons that have at least 3 trial repetitions of all the stimuli. The code below shows how we can store this information in an object called `sites_to_use` which we will use in our subsequent analyses.

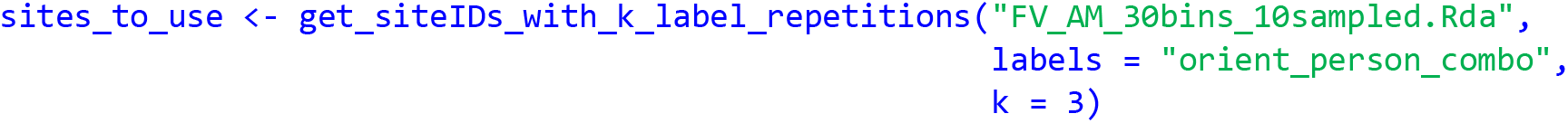

### Running the decoding analysis

Now that we have binned the data, and we have decided to use 3 cross-validation splits, we are ready to run the decoding analysis. To do this, we will first create a vector with the 25 strings that contain the names of label level for trials when left profile images were shown. We can do this by creating a vector of strings to restrict our decoding analysis to only decode trials when the left profile images were shown.

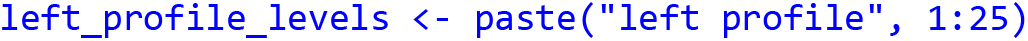

We can then run a decoding analysis by creating each of the 5 object types described in the Method section. For our first analysis we will use a:

1. `ds_basic` data source to create pseudo-populations of data. The arguments we pass to this constructor are: a) the name of the binned data file; b) the variable we want to decode (e.g., `“orient_person_combo”`); c) the number of cross-validation splits to use; d) a vector with the levels we want to use (i.e., the levels that start with “left profile”)^5^ and e) the site IDs for all the sites that have at least 3 label repetitions^6^.
2. `fp_zscore` feature preprocessor to normalize the data. This feature preprocessor ensures that neurons with higher firing rates do not dominate over neurons with lower firing rates. Feature pre-processes are put into a list which allow the analysis to contain more than one feature pre-processor, although we will only use one here.
3. `cl_max_correlation` classifier to make our predictions.
4. `rm_main_results` and `rm_confusion_matrix` result metrics to show our decoding accuracies. Result metrics are also put into a list which allows us to use two result metrics here.
5. `cv_standard` cross-validator to run the full decoding analysis. The arguments we pass to this constructor are: a) the data source; b) the classifier; c) the feature pre-processor; d) the result metrics; and e) a number specifying how many resample runs to use.

We set up these decoding objects using the code below:

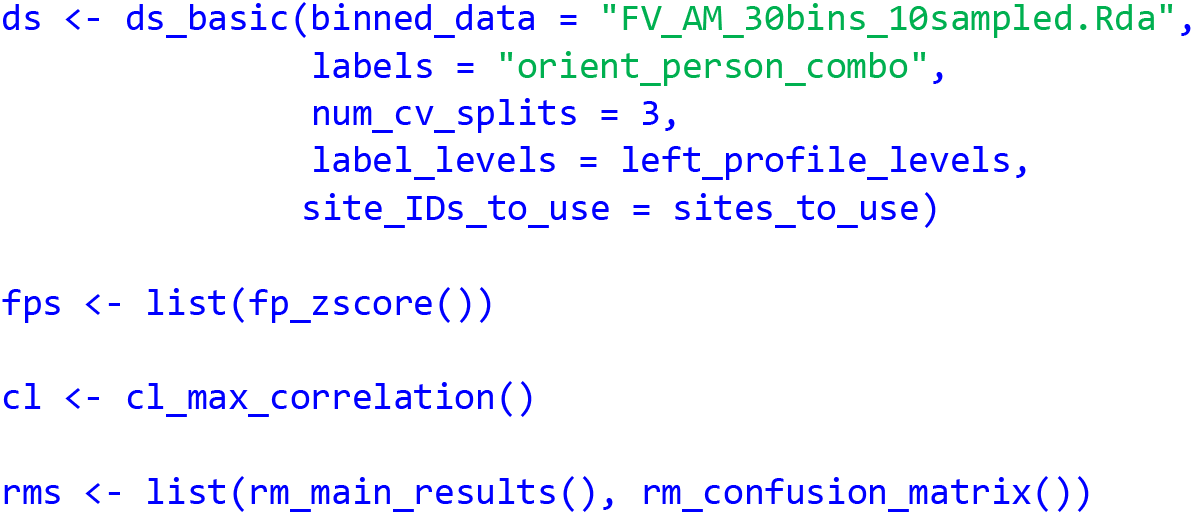

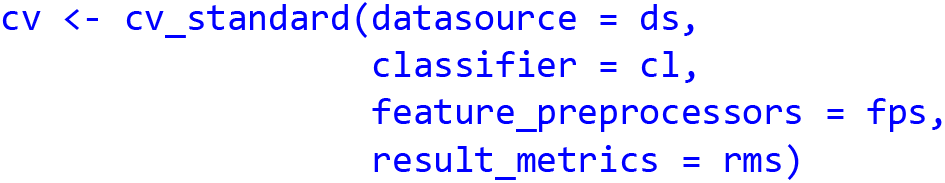

We can then run the decoding analysis using the cross-validators `run_decoding()` method as shown below where we can store the results from this analysis in a list that we usually name DECODING_RESULTS:

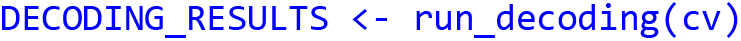

The `run_decoding()` function runs the decoding process in parallel, where the number of parallel cores can be specified by the optional `num_parallel_cores` argument to the `cv_standard` object (by default runs the code in parallel using half the cores that are available on the computer used for running the analysis)^7^.

### Plotting the results

The `DECODING_RESULTS` object created by running the `run_decoding()` function is a list that contains result metric objects which now hold the compiled results. Additionally, the `DECODING_RESULTS` object contains a list called `cross_validation_parameters` which stores the parameters that were used to generate the results. To see this for the `DECODING_RESULTS` object we created above, we can print out the names of the values stored in the `DECODING_RESULTS` object using:

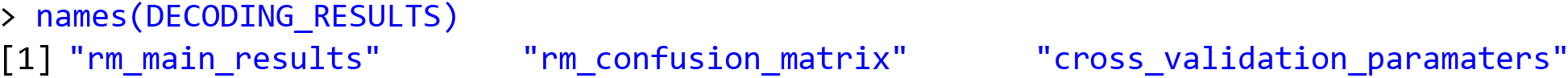

If we examine the `DECODING_RESULTS$cross_validation_parameters` list, we see it contains the objects used in the decoding analysis including `ds, fp, cl`, and `rm` objects, along with information about the analyses stored in the `parameter_df` data frame. If we examine the result metrics stored in the `DECODING_RESULTS` object, we see it holds the compiled results. In particular, the original empty result metrics that were passed to the cross-validator now hold the actual decoding results that were compiled from running `run_decoding(cv)` method. We can plot the results stored in these result metrics using their `plot()` functions.

The `rm_main_results` results metric `plot()` function creates a line plot of the decoding results as a function of time, as well as temporal cross decoding (TCD) plots. To create line plots, we use the `plot()` functions `type = “line”` argument. By default only the classification accuracy is plotted (i.e., the zero-one loss results), but we can also include normalized rank results, and the decision values on the plot by setting the `results_to_show = “all”` *a*rgument. The full call to the plot function then becomes:

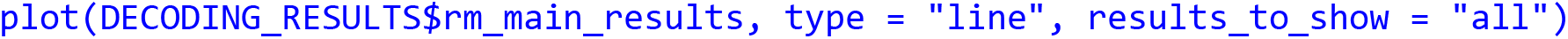

The results are show in Figure 3A. From looking at the results we see that the decoding accuracy rises above the chance level of 1/25 around 150 ms after stimulus onset, and the zero-one loss, normalized rank and raw decision value results look similar, which is often the case. If we want to plot a TCD plot, we set the `type = “TCD”`.

**Figure 3.**
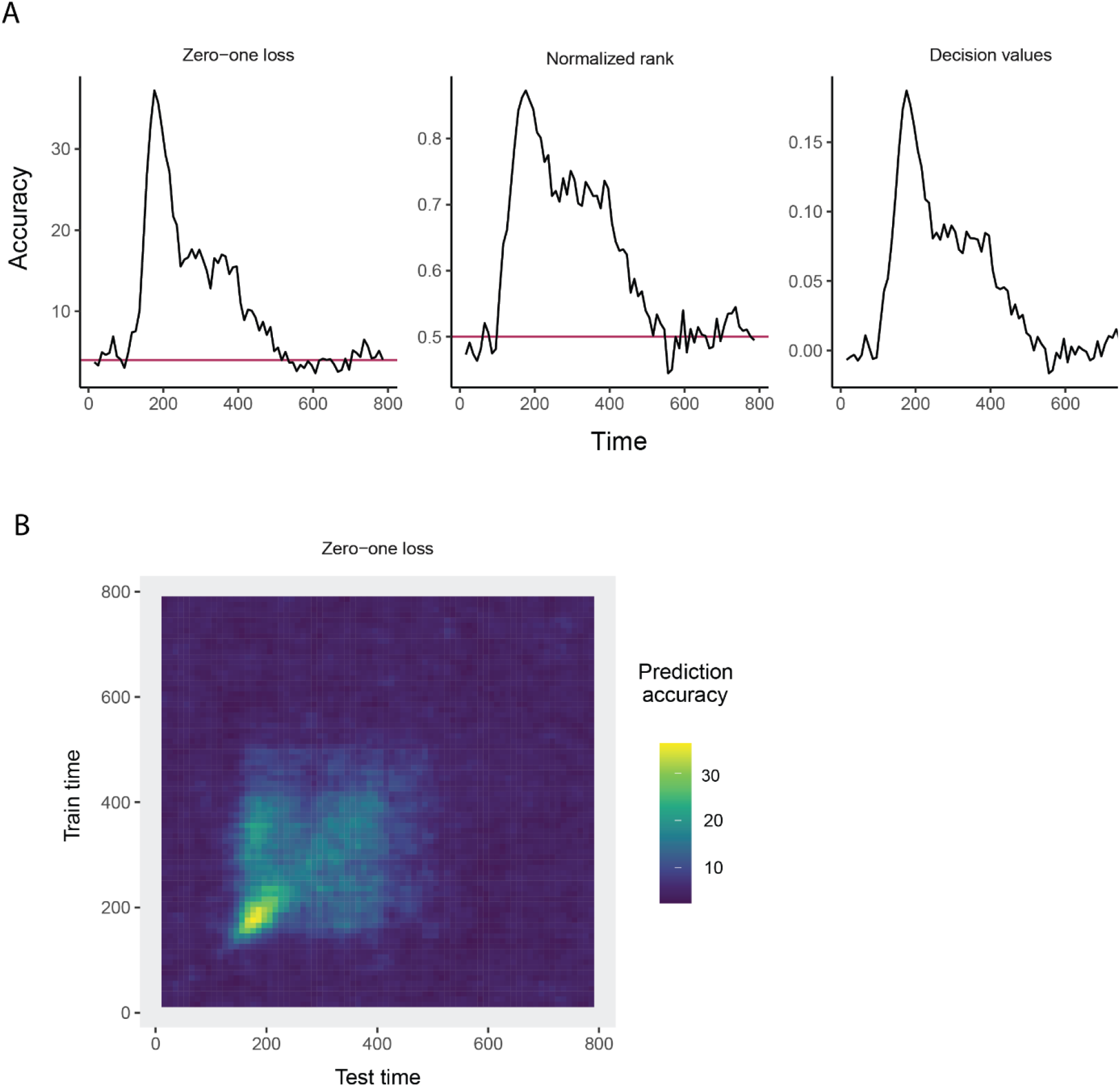
Results from decoding face identity (using only the left-profile face images) compiled by the rm_main_results object. **A**. A line plot showing the classification accuracy (zero-one loss), normalized rank, and decision value results based on running the `plot(rm_main_results, type = “line”)` function. As can be seen, the decoding accuracy increases above chance levels approximate 100 ms after stimulus onset. **B**. A temporal cross-decoding (TCD) plot based on using the `plot(rm_main_results, type = “TCD”)` function. The lack of a strong diagonal band on the plot indicates that information is not contained in a highly dynamic code.

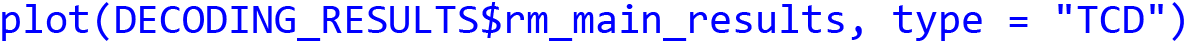

This TCD plot is in shown in Figure 3B. From looking at the TCD we do not see a strong diagonal yellow region in the plot which would indicate that the decoding accuracy is only high when training and testing the classifier at the same point in time. Thus, information appears to be contained in a stationary neural code where the same patterns of neural activity codes information at different points in time, rather than a dynamic population code where the patterns of neural activity that code information change in time.

The `rm_confusion_matrix` result metric `plot()` function allows one to view a sequence of confusion matrices for each time period that was decoded. Because we binned the data with a relatively small sampling interval of 10 ms, plotting a sequence of confusion matrices for all decoded time bins will be rather cluttered, so instead we will just plot the confusion matrix that starts around 200 ms after stimulus onset, which we can do by setting the argument `start_time_to_plot = 200`:

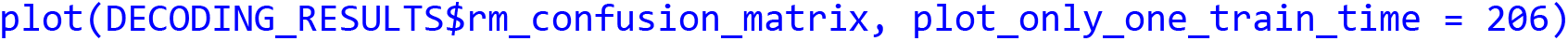

The results are shown in Figure 4. The y-axis on the plot shows the true class, and the x-axis on the plot shows the predicted class, and consequently diagonal elements on the plot are correct predictions. From looking at Figure 4, we see that some classes (i.e., face identities) were predicted more accurately than others, for example, the person 10 was correctly predicted about 60% of the time.

**Figure 4.**
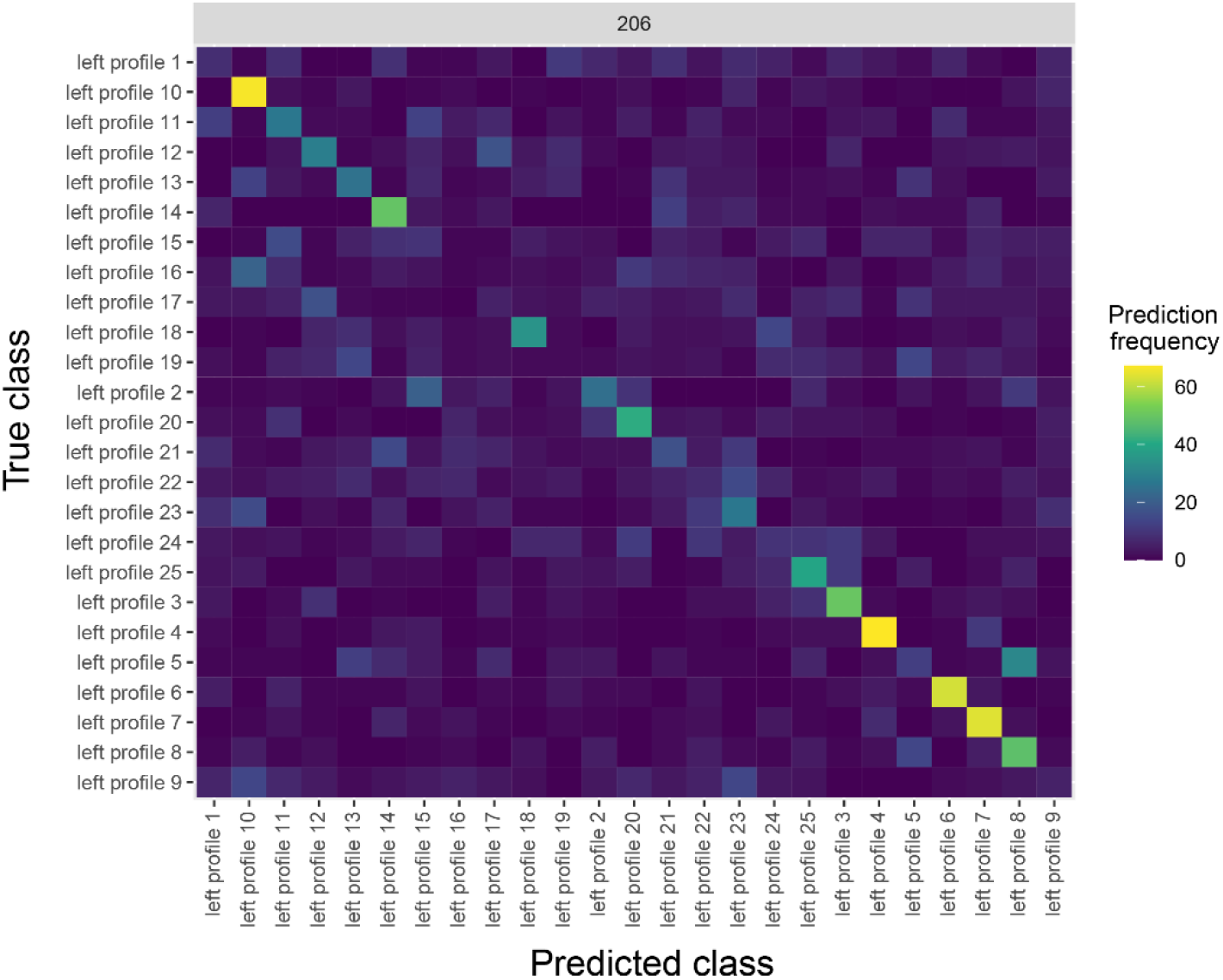
A Confusion matrix decoding face identity (using only the left-profile face images) compiled by the rm_confusion_matrix object. The confusion matrix is shown at 206 ms post stimulus onset using the function `plot(rm_confusion_matrix, plot_only_one_train_time = 206)`. Elements on the diagonal indicate a high level of correct predictions while elements off the diagonal indicate patterns of mistakes; for example, we that images of person 5 were fairly frequently classified as person 8.

### Saving and logging the results

After running an analysis, it is useful to save the results so that one can replot them at a later time, and compare the results to those from other analyses that have been run. Rather than using R’s `save()` function to save the results, it is useful to use the NeuroDecodeR’s `log_save_results()` function. This function not only saves the results, but it also creates a “manifest file” that keeps track of the parameters that were used in each decoding analysis that has run. This manifest file can then be used to search through all previous results that have been run, and to load previous results based on parameter values using the `log_load_results_from_params()` function.

To save the results using the `log_save_results()` function, we pass the `DECODING_RESULTS` list, along with a directory name where we would like to save the results. Additionally, we can also set the `result_name` argument to a string which will give a name to the results that can be used to reload the results using the `log_load_results_from_result_name()` function. For the analysis we ran above, we can save the results to a directory called “results” using the following:

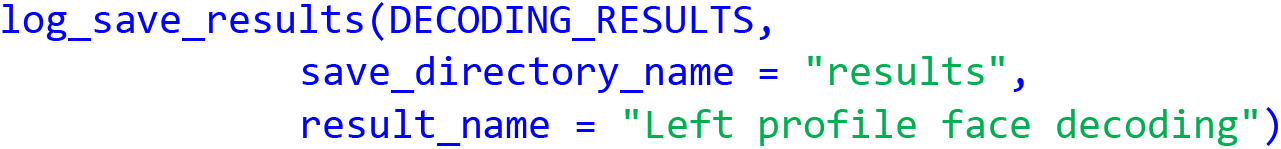

#### Decoding analysis 2: Testing generalization across head orientations

To further illustrate the capabilities of the NeuroDecodeR package, we now demonstrate how to run a “generalization analysis” where a classifier is trained on one set of conditions and then tested on a related set of conditions. In particular, we will train the classifier to distinguish between the 25 individuals using trials when left-profile face images were shown, as we did previously, but rather than testing the classifier on the same left-profile images (from different trials), we will instead test to see if the classifier can distinguish between the 25 individuals on trials when *right-profile* images were shown. If the classifier is able to classify the right-profile images at a high level of accuracy, this indicates that the neural representation contains information about face identity that is abstracted from the specific low-level visual features the classifier was trained on; i.e., anterior-medial face patch contains a representation of face identity that is invariant to the orientation of an individual’s head (Meyers et al., 2015).

Running this generalization analysis is very similar to running the basic decoding analysis we did above, however, we will use the `ds_generalization` data source rather than the `ds_basic` data source. The data, and all other NeuroDecodeR objects in this analysis will be the same as we used previously, which illustrates how the modular nature of the object in the NeuroDecodeR package allows one to easily run a range of analyses.

To create the `ds_generalization` data source, we will first create a vector of 25 strings that list all the right profile image names in the same order as the vector of left profile image names that we created previously. This can be done using:

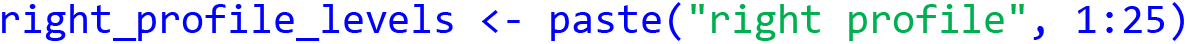

We can then create the `ds_generalization` datasource by specifying the same `binned_file_name, label_to_decode, num_cv_splits`, and `site_IDs_to_use`^8^ arguments as was done with the `ds_basic`, along with setting the `train_label_levels` to be the `left_profile_levels`, and the `test_label_levels` to be the `right_profile_levels`. Thus, the code for creating the `ds_generalization` is:

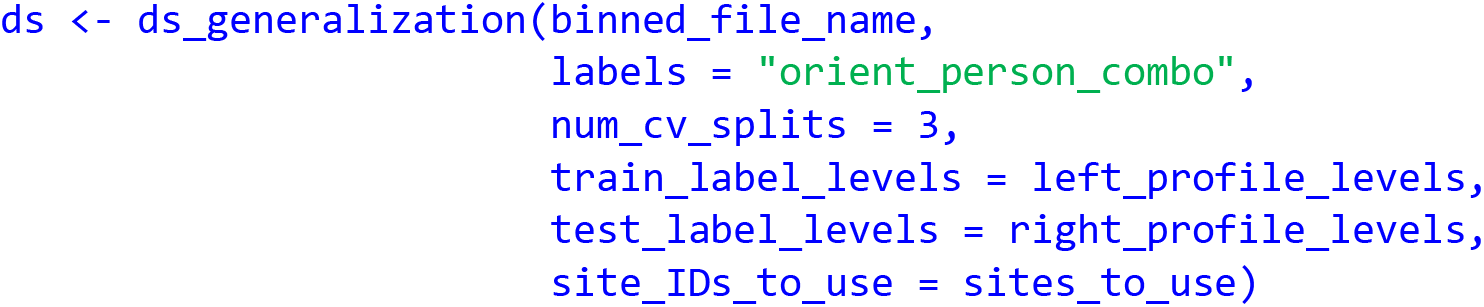

Now that the generalization data source has been created, we can create the other NeuroDecodeR objects, run the analysis, and save the results as was done before. To make the code run a little faster, we will omit using the rm_confusion_matrix, and we will omit creating the temporal cross decoding results by setting run_TCD = FALSE in the cross-validator.

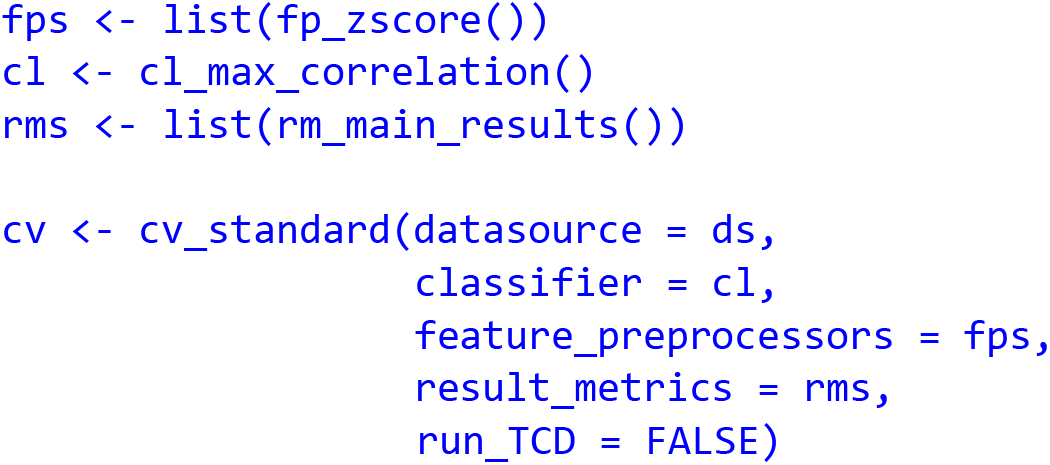

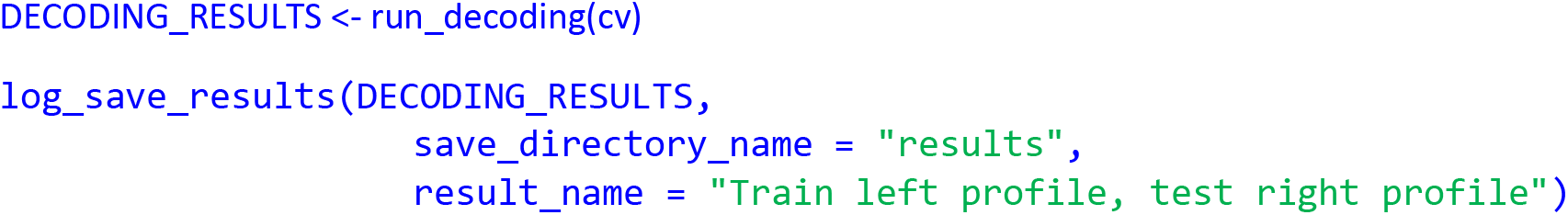

Once the analysis has finished running and we have saved the results, we can use `plot_main_results()` function to compare the results we created when training and testing the classifier on the left-profile images (that were created in the first analysis above) to the generalization results we just ran. To do this we will create a vector of strings that have the result names that we created when we saved the `DECODING_RESULT` objects from our two analyses. We will then pass the name of the directory holding the results, along with these result names to create a line plot that compares the results:

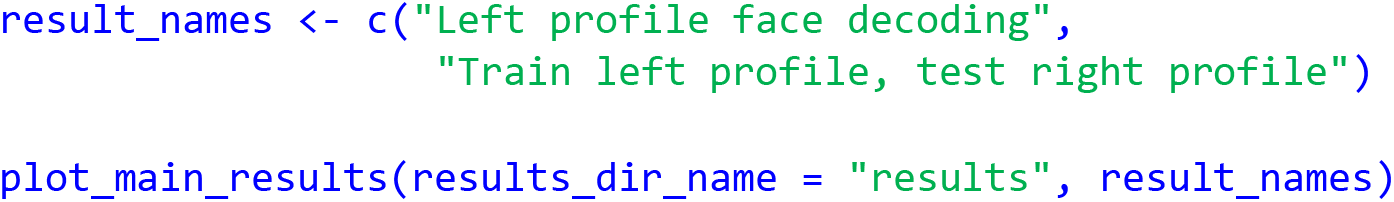

The resulting plot from running this code is shown in Figure 5. As can be seen, there is a similar level of decoding accuracies when the classifier is tested on the left and right profile images, which suggests that the neural representation of identity is highly invariant to the pose of the head, at least between left and right profile images. We encourage the reader to experiment with assessing the invariance across other head orientations, such as training on the front facing images and testing on the profile images^9^.

**Figure 5.**
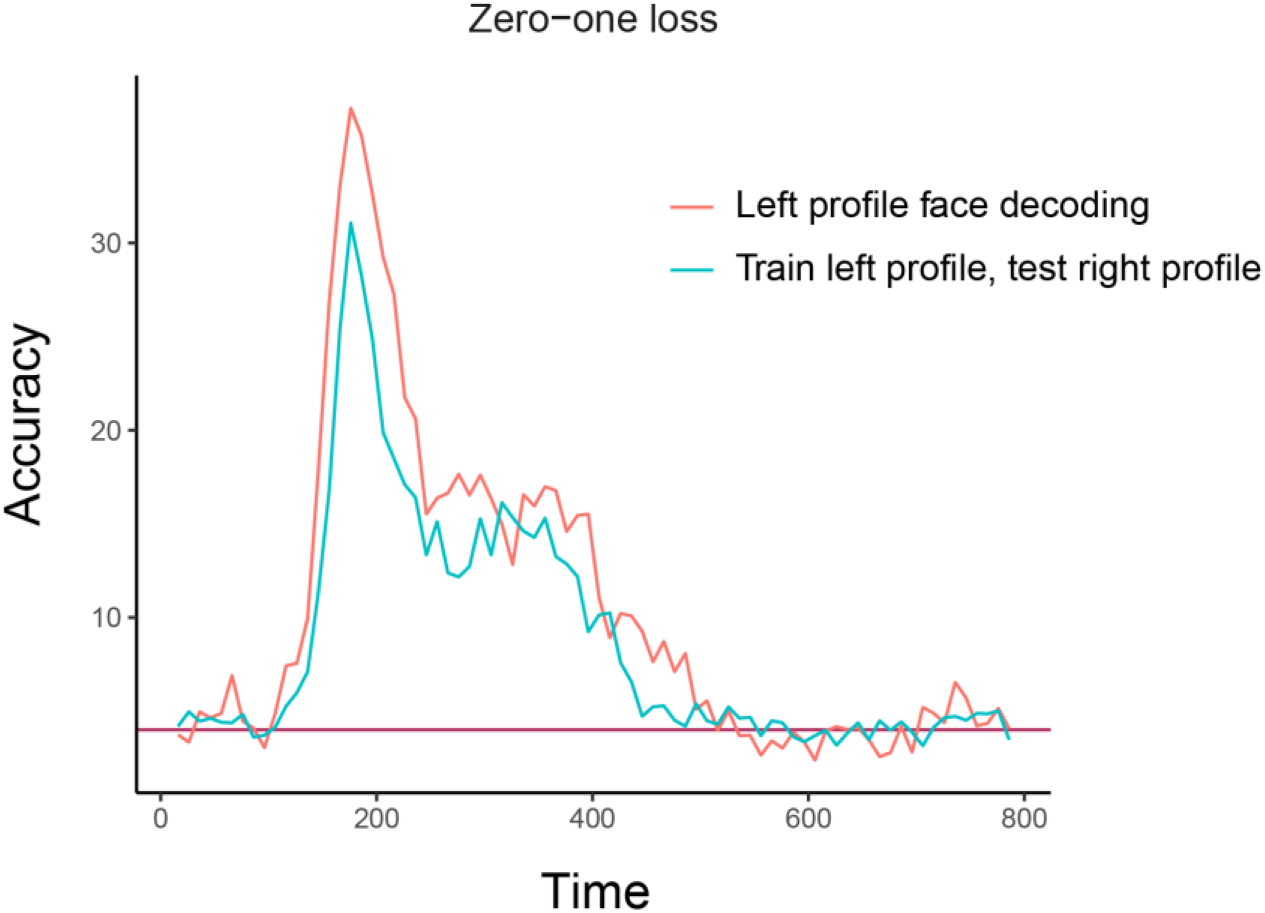
Comparison of basic identity decoding results to generalization analysis results. The plot shows the results from classifying the 25 individual face identities using a basic decoding analysis where the classifier was trained and tested using on left profile face images (red trace) to generalization analysis results where the classifier was trained on left profile images and tested on right profile images (cyan trace). As can be seen, the classification accuracies are fairly similar for the basic and generalization analysis indicating that face identity information in brain region AM is contained in a code that is highly invariance to the pose of the head across left and right profile images.

#### Piping together NeuroDecodeR objects

A popular way to write data analysis code in R is to use the pipe operator to string together a sequence of data analysis functions. The NeuroDecodeR package also supports the pipe operator to create a sequence of functions that are needed to run a decoding analysis. This can be done by:

1. Starting with a binned data file name
2. piping it to a data source
3. piping together feature-preprocessors, a classifier, and result metrics
4. piping this to a cross-validator.

The code below illustrates how to run the same basic analysis as our first analysis above using the pipe operator. For the sake of novelty, we will decode the right profile images, and we plot the results from all the analyses we have run (see supplemental Figure 1).

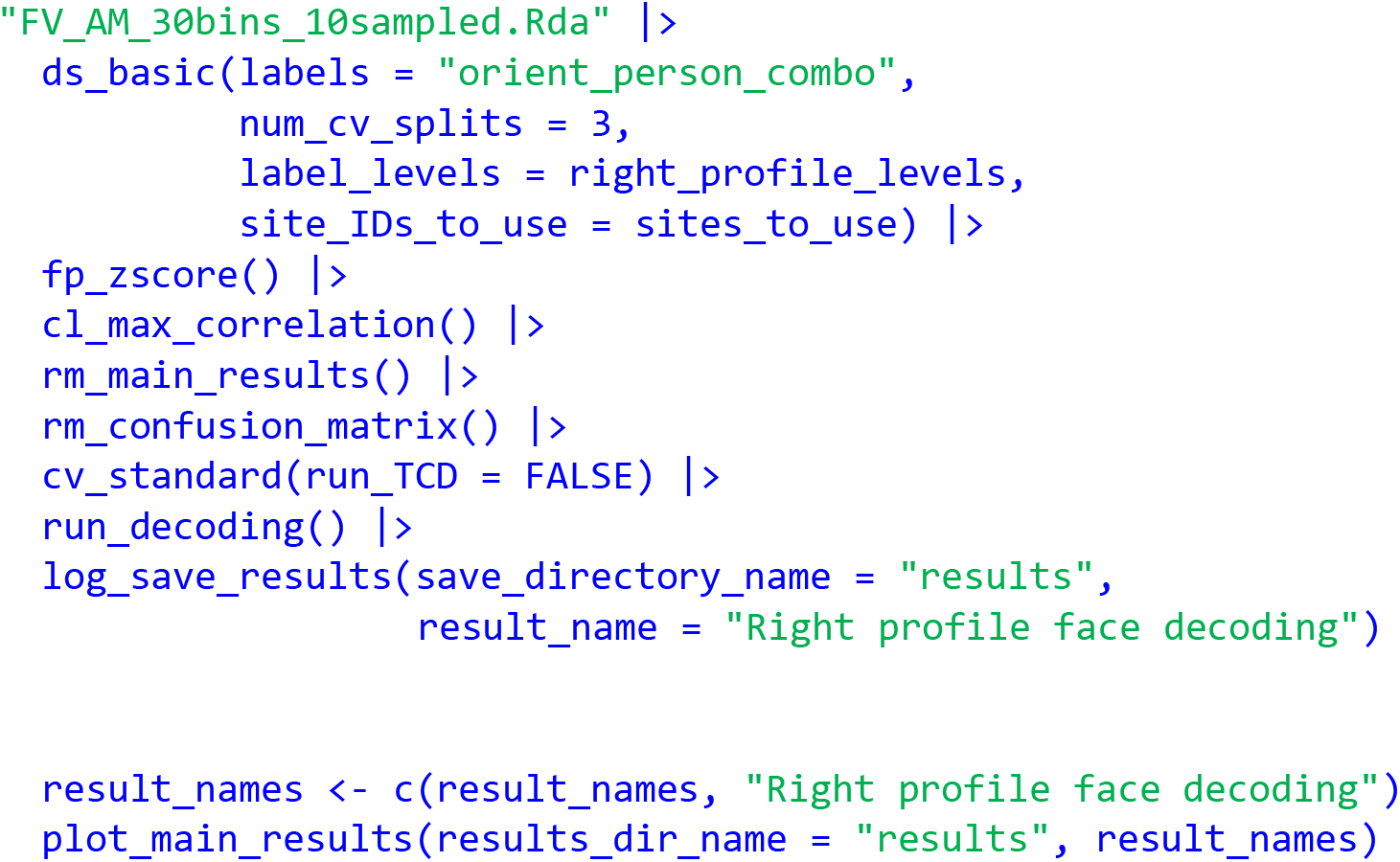

## Discussion

In this paper, we introduced the NeuroDecodeR package, described the design and data formats used by the package, and gave examples of how to run a basic decoding and generalization analysis. We also described the benefits of using the package, including the fact that the modular design of the package makes it easy to run a range of different decoding analyses, the decoding code is parallelized which speeds up the run time of analyses, and a system that makes it easy to save and manage decoding results. Additionally, using the R programming language to run decoding analyses has several benefits including that it is a free/open source language, and that R has a strong ecosystem for creating reproducible analyses. In particular, we highly recommend that users of the NeuroDecodeR package take advantage of these reproducible features in R by creating R Markdown documents that contain the code used for each analysis so that it is easy to recreate any analysis run^10^.

The description of the NeuroDecodeR package in this paper is based on the package’s initial release, however, we anticipate continuing to develop and extend the package. A few directions we are interested in extending the package include adding additional NeuroDecodeR objects (e.g., additional data sources and classifiers), and writing code that can give estimates of the memory usage and runtime that particular analysis will take which will enable users to choose decoding parameters based on the computing system they are using for their analyses. For example, users could bin their data at a higher temporal resolution if they are running their analyses on a more powerful computer. We also plan to create a shiny app for the package that will allow users to generate R Markdown documents that contain all the code needed to run an analysis by clicking buttons on a graphical user interface. This “NeuroShiny” app will further shorten the time it takes to run decoding analyses, and will enable Neuroscientists with little programming experience to run reproducible decoding analyses on their data. Finally, we anticipate creating other types of analyses that use the same binned and raster formats described in this paper, which will allow researchers to easily run a whole range of different types of analyses once they have put their data into the proper format.

In summary, we believe that the NeuroDecodeR package will be of great benefit to neuroscientists who are interested in using decoding to analyze their data, and should also offer an entry point for Statisticians who are familiar with the R programming language to become involved in analyzing neural data. We hope that this package will lead to new insights into how the brain processes information and will help to speed up the pace of discovery in neuroscience.

## Acknowledgements

We would like to thank Tomaso Poggio for his continued support, and Xinzhu Fang, Sukolta Rithichoo, Xinyao Lu, Boming Zhang, Yongyi Peng, Elisa Loy, and Jenny Pan, for helping to test the NeuroDecodeR package. We would also like to thank Doris Tsao, Winrich Friewald, Christos Constantinidis, Robert Desimone and Ying Zhang for their willingness to share data they collected.

## Supplemental material

**Supplemental figure 1.**
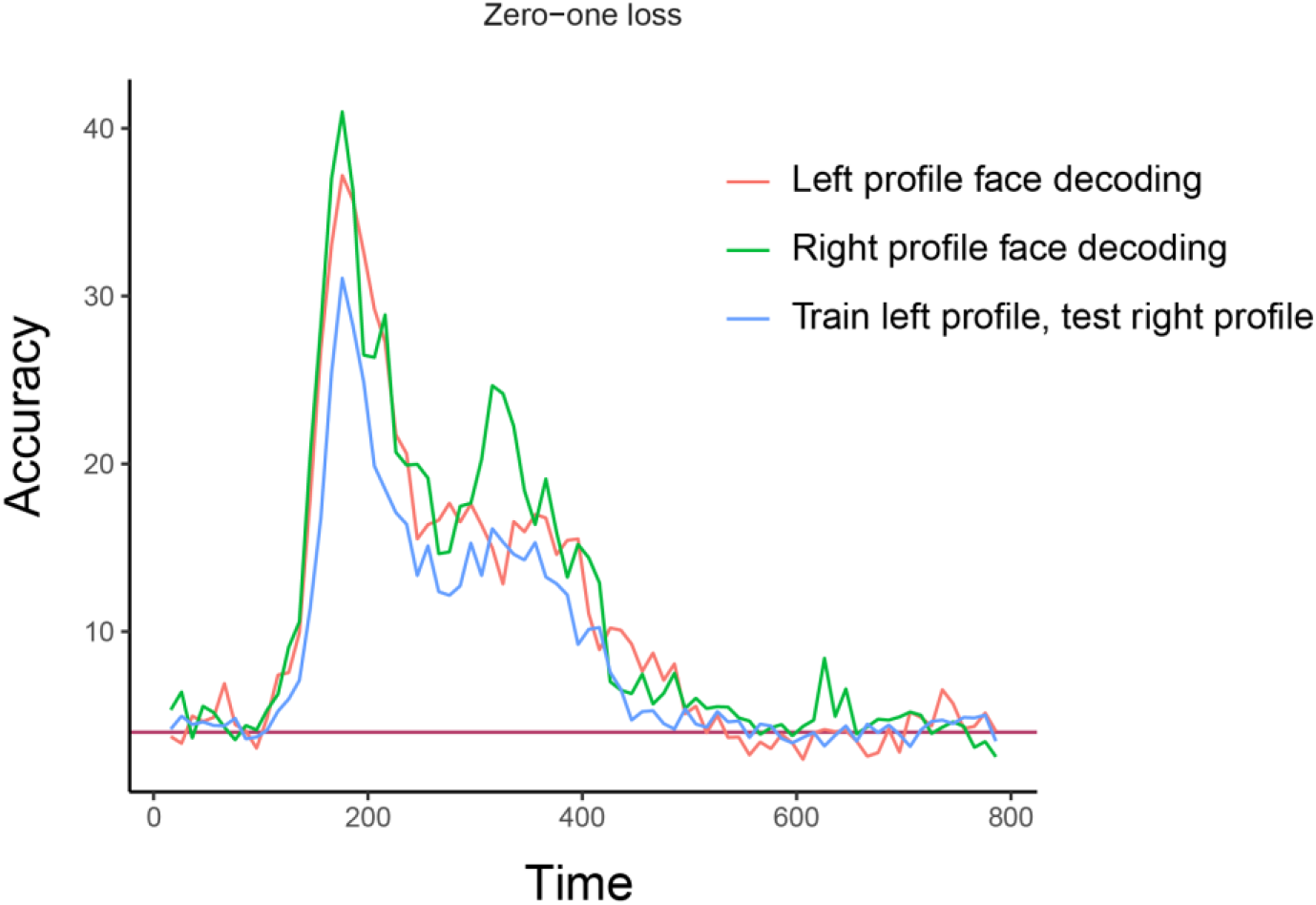
Comparison of basic identity decoding results for left and right profile images, to generalization analysis results. The plot shows the results from classifying the 25 individual face identities using a basic decoding analysis where the classifier was trained and tested using on left profile face images (red trace), or on right profile images (green trace) to generalization analysis results where the classifier was trained on left profile images and tested on right profile images (blue trace); i.e., these are the same results as Figure 5 except the basic decoding results of training and testing on the right profile image have been added to the plot. Again, we see the classification accuracies are fairly similar for the basic and generalization analysis indicating that face identity information in brain region AM is contained in a code that is highly invariance to the pose of the head across left and right profile images.

**Supplemental Table S1.**
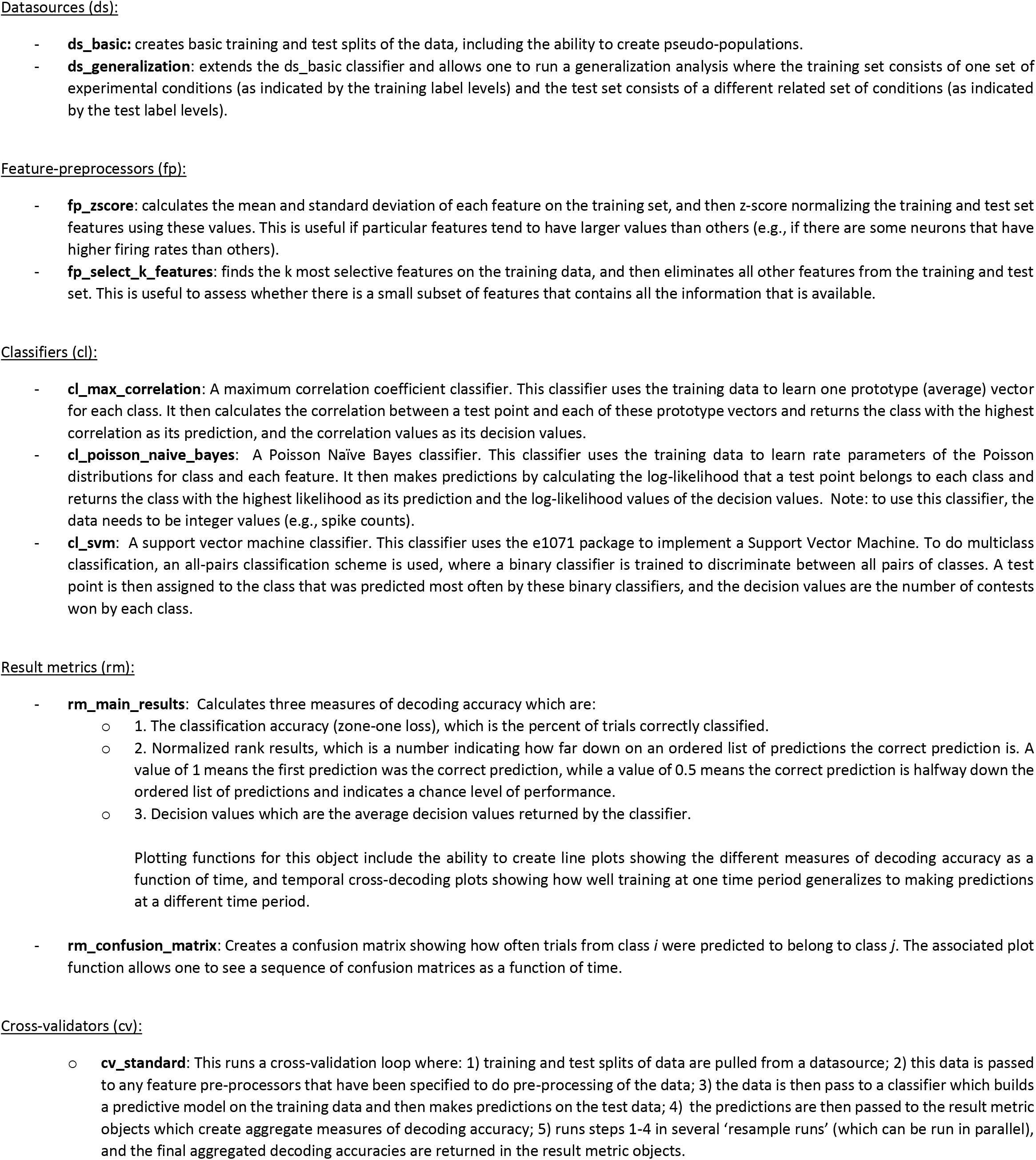
More detailed description of implementations of NeuroDecodeR abstract objects that come with the NeuroDecodeR package. The package also includes a number of additional functions that are useful for plotting and processing the data, and for saving and loading results. For more details on these objects, see the online documentation at: https://emeyers.github.io/NeuroDecodeR/reference/index.html

## Footnotes

1 Experimenting with different classification algorithms is useful for gaining insight into the ‘neural code’ and for assessing the maximum amount of information that could be extracted by a sequence of downstream areas (Glaser et al., 2020; E. M. Meyers et al., 2015).

2 BioRxiv does not allow images of faces to be displayed so we can not show examples of these stimuli. However, examples of the stimuli can be seen in Figure 1C of Meyers et al (2015).

3 There are also a few additional optional arguments to control the binning process. To see all the arguments, one can view the function documentation by typing ? `create_binned_data()`.

4 Using only 3 cross-validation splits is generally considered a very low number for decoding analyses, and we would prefer to use 6 or more splits. However, given that the AM brain region is highly selective for faces, as you will see below, this analysis still gives useful results.

5 If the `label_levels` argument is not specified, all available label levels will be used in decoding, which would be 200 stimuli in this case (i.e., 25 individuals from 8 head orientations).

6 If this argument is not specified, then all sites that have at least 3 repetitions for the label_levels specified will be used. This is slightly different than the sites we have selected since we are only using sites that have at least 3 repetitions *for all 200 stimuli*, rather than at least 3 repetitions for only the left-profile images. The reason we are using only sites with 3 repetitions of *all* stimuli, is so that we can make a fair comparison to decoding right profile images in the second analyses that is described below.

7 The code took a little over 8 minutes on the system we tested it on. However, the run time will obviously depend on the specifications of the computer you use for the analysis. If you would like the code to run faster, you can try binning the code using a larger sampling interval of 30 ms, which cut down the run time on our computer to be a little over 2 minutes.

8 It is possible that different sites have different numbers of left and right profile trials, which would make different sites available for decoding left profile and right profile images. By setting the `site_IDs_to_use` in both the basic and generalization analyses using only sites that have at least 3 repetitions for all head orientation images, we guarantee that the same sites will be used in our basic decoding analysis (train and test left) and in our generalization analysis (train left test right). This allows a fair comparison of these analyses since differences in decoding accuracies cannot be due to different neurons being used.

9 Spoiler, decoding accuracy on the frontal images is higher, and there is less invariance generalizing from the frontal images to the profile images.

10 The fact that the decoding parameters are saved in the `DECODING_RESULTS` also allows one to easily verify the parameters that were used in an analysis. However the use of R Markdown to save the code as a pdf allows one to directly store the code used which can make it easier to read.

